# RAISS: Robust and Accurate imputation from Summary Statistics

**DOI:** 10.1101/502880

**Authors:** Hanna Julienne, Huwenbo Shi, Bogdan Pasaniuc, Hugues Aschard

## Abstract

**Motivation:** Multi-trait analyses using public summary statistics from genome-wide association studies (GWAS) are becoming increasingly popular. A constraint of multi-trait methods is that they require complete summary data for all traits. While methods for the imputation of summary statistics exist, they lack precision for genetic variants with small effect size. This is benign for univariate analyses where only variants with large effect size are selected a posteriori. However, it can lead to strong p-value inflation in multi-trait testing. Here we present a new approach that improve the existing imputation methods and reach a precision suitable for multi-trait analyses.

**Results:** We fine-tuned parameters to obtain a very high accuracy imputation from summary statistics. We demonstrate this accuracy for small size-effect variants on real data of 28 GWAS. We implemented the resulting methodology in a python package specially designed to efficiently impute multiple GWAS in parallel.

**Availability:** The python package is available at: https://gitlab.pasteur.fr/statistical-genetics/raiss, its accompanying documentation is accessible here http://statistical-genetics.pages.pasteur.fr/raiss/.

## 1 Introduction

By solving practical and ethical challenges, public summary statistics has become a gold entry point for the study of complex traits (B. Pasaniuc & Price, 2017). In the past years, multi-trait methods using summary statistics have attracted much scientific attention and yields many applications including e.g. multi-trait testing (Liu & Lin, 2018; Turley et al., 2018) or correction for pleiotropy in mendelian randomization (Verbanck, Chen, Neale, & Do, 2018). Most multi-trait methods are only applicable to single nucleotide polymorphisms (SNPs) with complete data for the traits of interest, and imputation of missing statistics is mandatory in many real data analyses. However, in the multi-traits context, imputation must reach a very high level of accuracy, even SNPs with moderate effect size, to avoid false association signal. Existing solutions do not achieve this level of accuracy (see Supplementary Fig.1). The imputation must also be time efficient so computation does not become a bottleneck in pre-processing many traits.

We improved an existing imputation solution (Bogdan Pasaniuc et al., 2014) on two key points to make it suitable for multi-trait applications: 1) we optimize a hyper-parameters through a systematic space search. 2) We designed the python package RAISS (Robust and Accurate imputation from Summary Statistics) so multiple traits can be imputed in parallel.

## 2 Methods

### 2.1 Statistical model

The statistical model used in RAISS is similar to the one described in (Lee, Bigdeli, Riley, Fanous, & Bacanu, 2013). Summary statistics are given as Z-scores. The idea behind summary statistics imputation is to leverage linkage disequilibrium (LD) to compute Z-scores of unknown SNPs from neighboring typed SNPs:

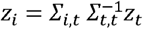

Where is the vector of SNPs to impute, is the vector of typed SNPs and Σ is linkage disequilibrium matrix between SNPs.

### 2.2 Ensuring correct inversion of

Neighboring SNPs are highly correlated variables which makes the inversion of prone to numerical instabilities. We invert with the Moore-Penrose pseudo inverse. To ensure numerical stability, we applied a very stringent pruning of small eigen-values –*i.e*. eigen values below a given threshold are set to zero in the computation of pseudo inverse (*rcond* parameter in the scipy.linalg.pinv function).

### 2.3 RAISS pipeline and computation time optimization

#### 2.3.1 Precomputation of linkage disequilibrium

We derived LD using individuals of European ancestry from the 1000 genome panel (Genomes Project et al., 2012) (see RAISS documentation). To avoid repeated estimation of LD when imputing statistics for multiple GWAS, RAISS precompute pairwise linkage disequilibrium between SNPs present in the reference panel (see Supplementary data and Supplementary Figures 2 and 3).

#### 2.3.2 Command line tool for chromosome imputation

The simplest access to the imputation function in RAISS is the shell command *raiss* (accessible in a terminal after installing the package). This command impute the summary statistics for one trait and one chromosome and filter the results according to the predicted (see Supplementary Fig.2 and RAISS documentation).

## 3 Results

We tested RAISS performances using the following procedure:

For a chromosome and a trait,

1. Remove randomly 5000 SNPs in the Z-score file
2. Impute these 5000 SNPs
3. Set imputation hyper-parameters
4. Compute the correlation between the real Zscores and the imputed Zscores

### 3.1 Effect of hyper-parameters

We ran the above validation procedure on chromosome 22 for a Height GWAS (Wood et al., 2014). We varied the stringency in pruning small eigen-value from 10e-15 (default value in scipy) to 0.1, and the predicted imputation *R*^2^ filtering threshold from 0.1 to 0.9.

The pruning threshold for small eigen-value turns out to be the most important hyper-parameter to ensure a good correspondence between typed and imputed Z-scores. The imputation quality concomitantly increases with the threshold to reach a high correlation of 0.96 (see Figure 1). Filtering imputed SNPs by their predicted R^2^ improves only slightly the imputation accuracy. Moreover, if the R^2^ threshold is set too high most of the imputed SNP would be filtered (see Supplementary Fig.4 and Supplementary data).

**Figure 1.**
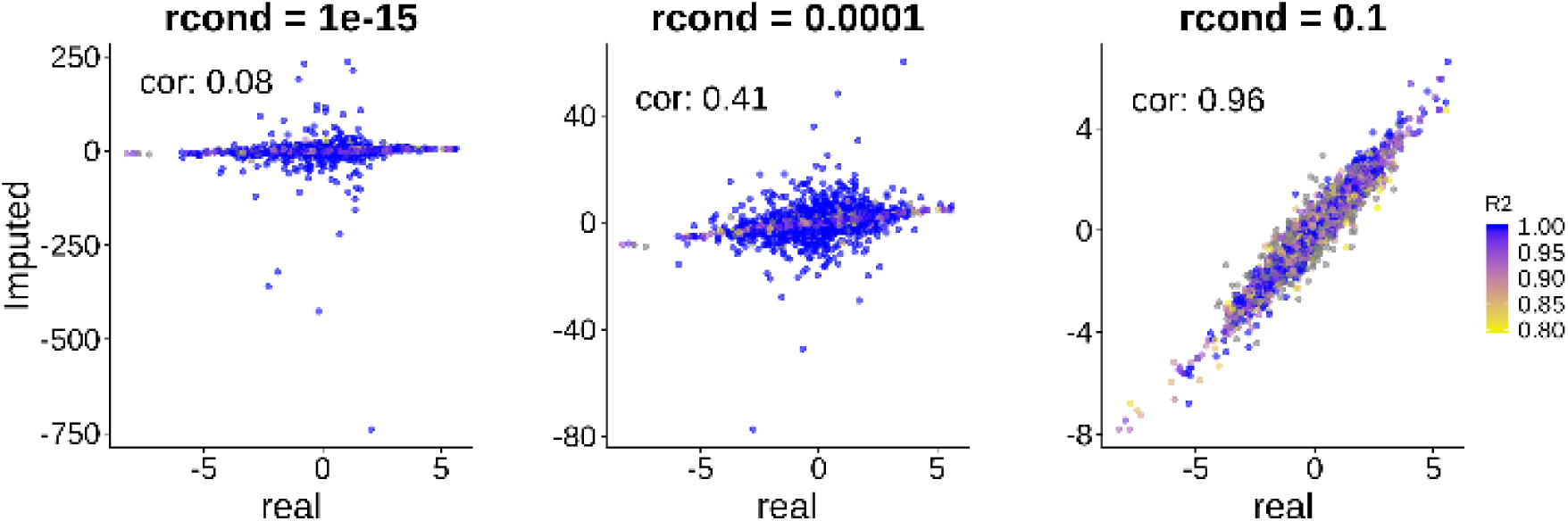
Real versus Imputed zscores for three different eigen value tresholds after filtering point with a R^2^ bellow 0.8.

### 3.2 Performance on a large panel of traits

To further assess the relevance of the hyper-parameters defined using the height GWAS data (*rcond* = 0.1 and *R*^2^ >0.6), we applied the final procedure for the analysis of 28 GWAS (see Supplementary table 1). The correlation between real and imputed Z-scores varied from 0.9 to 0.97 (see Supplementary table 2) dramatically increasing performances as compared to existing approach. We used imputed summary statistics from RAISS as input for a multivariate test method currently available at http://jass.pasteur.fr/index.html and we did not observe any p-value inflation as measured by the genomic control coefficient (see Supplementary Fig. 5).

## 4 Conclusion

We implemented an efficient tool allowing for the imputation of multiple summary statistics in parallel. We demonstrate a greatly improved accuracy for small size-effect variants in the real data analysis of 28 GWAS. Thus, the RAISS package an appropriate level of confidence that makes it suited for the imputation of summary statistic for various multi-trait analyses.

