## Supplementary data for "RAISS: Robust and Accurate imputation from Summary Statistics"

Table of content

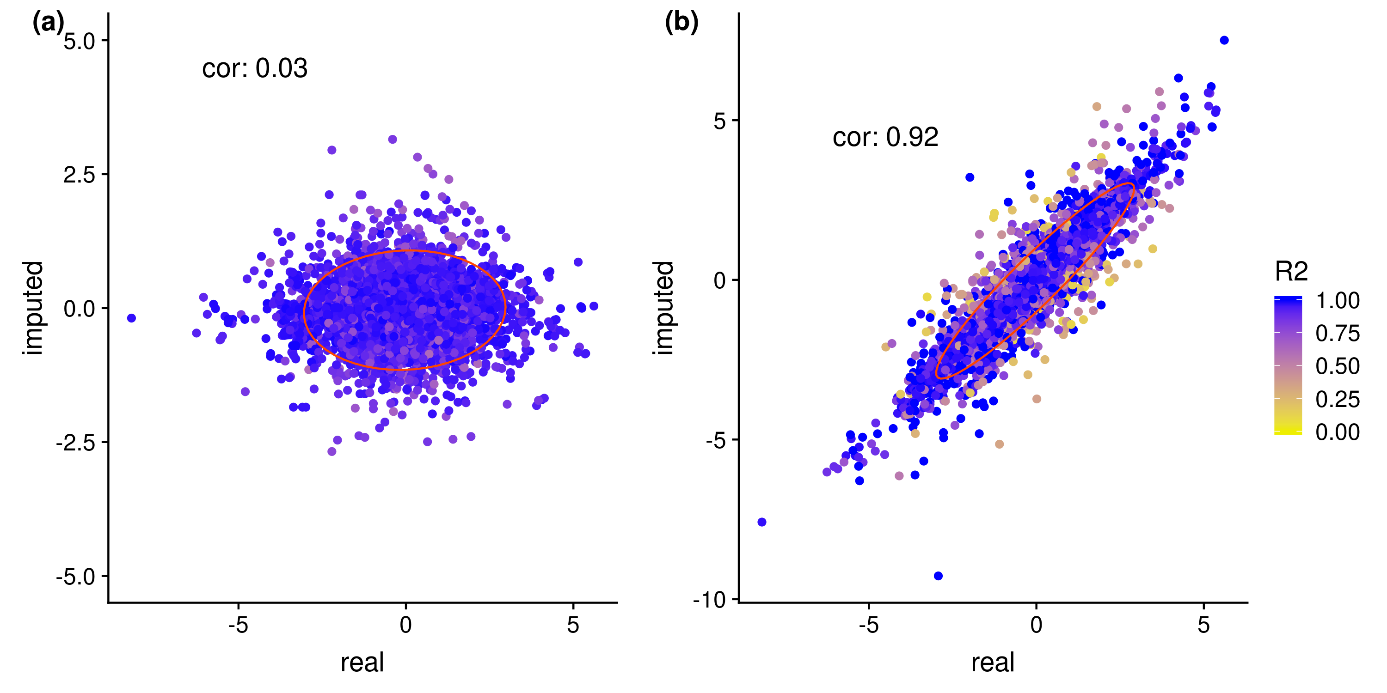

### Comparison of imputation accuracy between IMPG and RAISS

Supplementary Fig. 1 Imputation accuracy for small size-effect SNPs. Real values versus imputed values for default parameters (window_size= 1000kbp, lambda=0.1) for (a) IMPG and (b) RAISS.

### RAISS pipeline overview

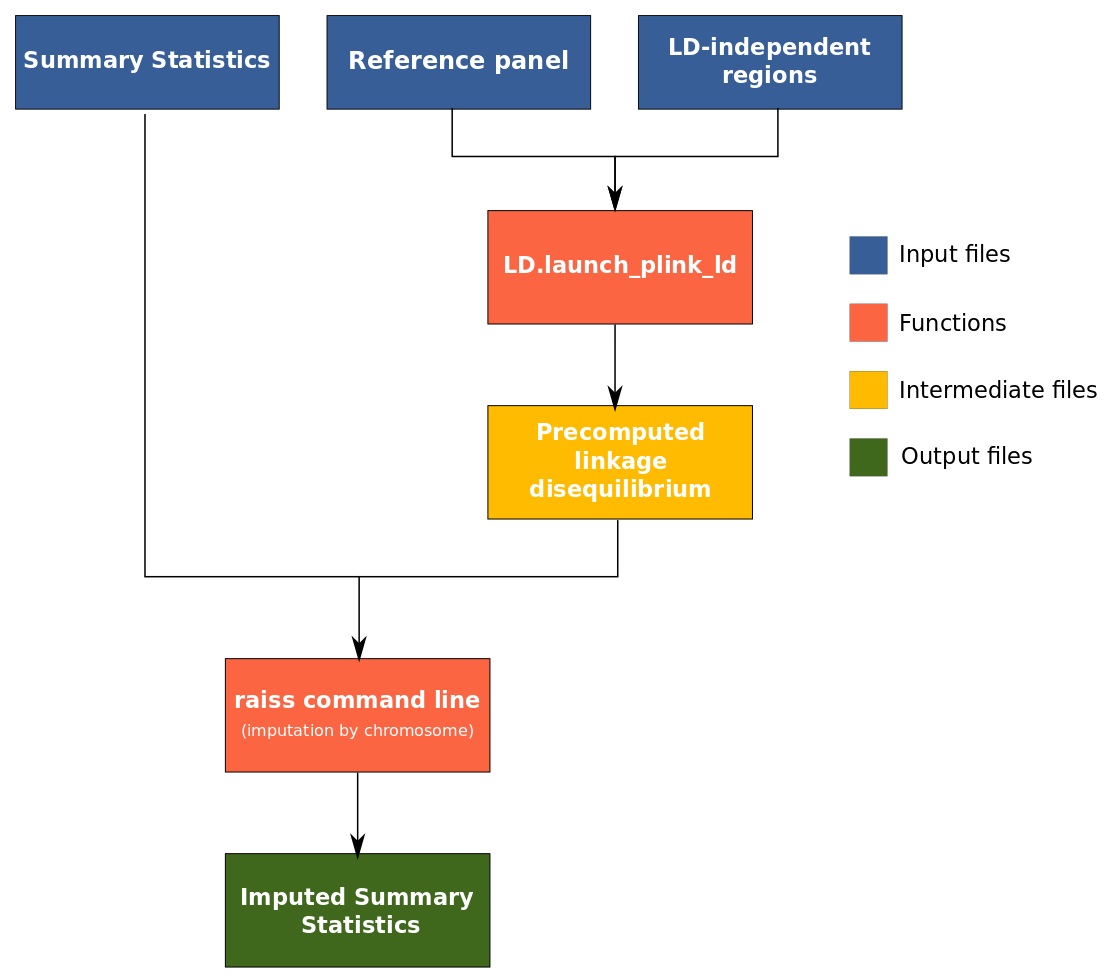

Supplementary Fig. 2 RAISS pipeline for imputation

### LD precomputation

To limit the size of the LD intermediate files, we precompute LD only between pairs of SNPs inside the same LD independent region (Berisa & Pickrell, 2016) (could be replaced by arbitrary region of ~1Mb) and save it as a tabular file. Note that this represents a sizeable amount of data when using the default LD block definition (a few dozen of Gb). Once computed, these files are accessed during the imputation of all traits. This solution avoid to re-compute the LD for each trait and hence use storage space to save computation time.

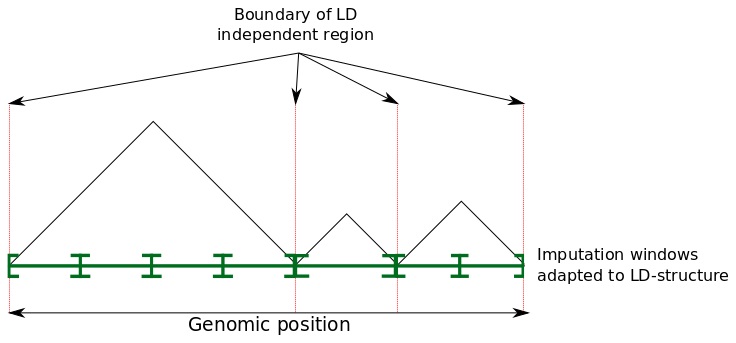

Supplementary Fig. 3 Position of imputation windows relatively to LD-independent region. In Raiss implementation we adapt imputation windows to LD regions.

### Choice of the $\boldsymbol{R}^{\mathbf{2}}$ threshold

Filtering imputed SNPs by their predicted $R^{2}$improves slightly the imputation accuracy. However, if the $R^{2}$ threshold is set too high most of the imputed SNP would be filtered. On the 28 traits used (see section 3.2 of the main document), we computed the fraction of the imputed SNPs filtered out (Supplementary figure 4 (a)) and the correlation between the imputed and typed Z-scores versus the imputation $R^{2}$ threshold (Supplementary figure 4 (b)). A $R^{2}$ threshold set at 0.6 is a good trade-off enabling to keep ~75% of the imputed SNPs while improving imputation accuracy compared to no filtering. Another possible value would be 0.8 which is optimal for imputation accuracy.

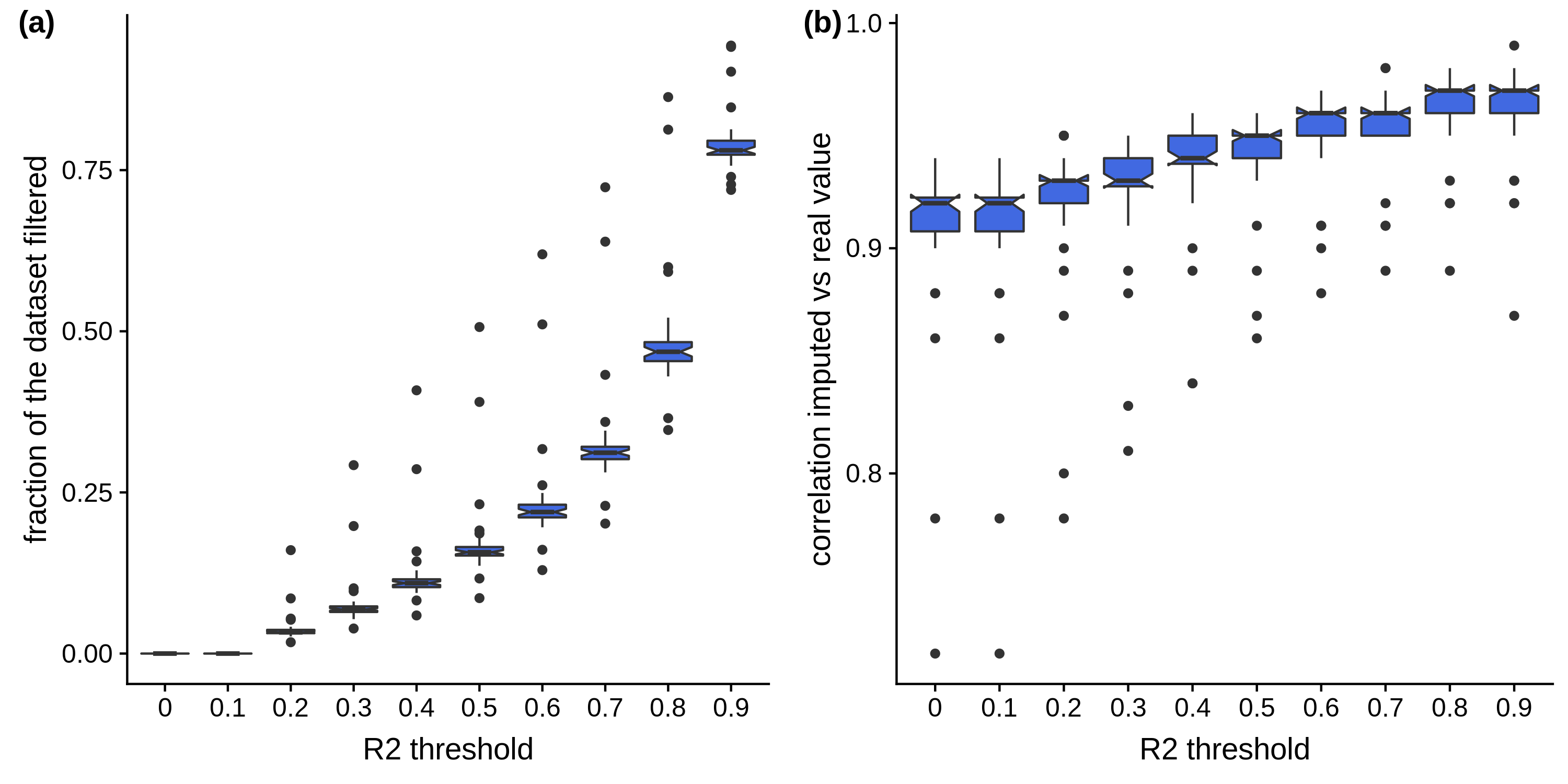

Supplementary Fig. 4 For the 28 traits tested (a) fraction of the dataset filtered versus the $R^{2}$ threshold. (b) correlation between the imputed and typed Z-scores versus the imputation threshold.

### Absence of inflation after imputation in an example of multi-trait testing

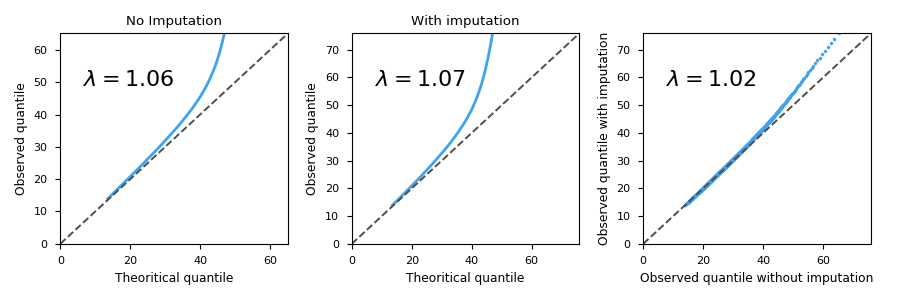

Supplementary Fig. 5 Effect of imputation of missing z-scores on the test statistic. All 28 traits used in the studies where tested conjointly. The left column is the empirical quantile of the statistic versus the theoretical quantile before imputation. The middle column is the same after imputation. The right column is the empirical quantile of the statistic before imputation versus the same quantity after imputation.
